# Optimal CXCR5 Expression during Tfh Maturation Involves the Bhlhe40-Pou2af1 Axis

**DOI:** 10.1101/2024.05.16.594397

**Authors:** Xiaoliang Zhu, Xi Chen, Yaqiang Cao, Chengyu Liu, Gangqing Hu, Sundar Ganesan, Tibor Z. Veres, Difeng Fang, Shuai Liu, Hyunwoo Chung, Ronald N. Germain, Pamela L. Schwartzberg, Keji Zhao, Jinfang Zhu

## Abstract

The pair of transcription factors Bcl6-Blimp1 is well-known for follicular T helper (Tfh) cell fate determination, however, the mechanism(s) for Bcl6-independent regulation of CXCR5 during Tfh migration into germinal center (GC) is still unclear. In this study, we uncovered another pair of transcription factors, Bhlhe40-Pou2af1, that regulates CXCR5 expression. Pou2af1 was specifically expressed in Tfh cells whereas Bhlhe40 expression was found high in non-Tfh cells. Pou2af1 promoted Tfh formation and migration into GC by upregulating CXCR5 but not Bcl6, while Bhlhe40 repressed this process by inhibiting Pou2af1 expression. RNA-Seq analysis of antigen-specific Tfh cells generated in vivo confirmed the role of Bhlhe40-Pou2af1 axis in regulating optimal CXCR5 expression. Thus, the regulation of CXCR5 expression and migration of Tfh cells into GC involves a transcriptional regulatory circuit consisting of Bhlhe40 and Pou2af1, which operates independent of the Bcl6-Blimp1 circuit that determines the Tfh cell fate.

---

Tfh cells, as one subset of T helper cells, play a critical role during humoral immune response by helping B cells in antibody production^1^. Similar to other effector CD4 T (Teff, e.g., Th1/Th2/Th17) cells, Tfh cells differentiate from naïve CD4 T cells, and their differentiation is tightly regulated by TCR signaling and cytokine milieu. Bcl6 has been identified as the master transcription factor determining the Tfh cell fate, establishing Tfh cells as another distinct lineage of Teff cells^2, 3, 4^.

During the early phase of naïve CD4 T cell differentiating into pre-Tfh cells induced by antigen presented by dendritic cells (DCs), Bcl6 upregulation determines Tfh cell fate whereas Blimp1 upregulation results in other Teff cells^3^. Strength of TCR signaling might play an important role during Tfh and non-Tfh fate determination^5^. For the second priming, pre-Tfh cells contact with congenic B cells and receive additional signals from B cells for further development, via ICOS-ICOSL^6^ and SLAM family members^7^. During the late phase of Tfh cell differentiation, further upregulation of the chemokine receptor CXCR5 and downregulation of the T cell zone–homing chemokine receptor CCR7 are critical for guiding Tfh cell migration into germinal center (GC) to become GC-Tfh^8, 9, 10^. In GCs, GC-Tfh exert their functions by helping B cells go through affinity maturation to become high affinity antibody-secreting plasma cells or long lived memory B-cells, a process that is important for host’s defense against various pathogens^11^.

The location of Tfh cells determines their functionality^12, 13^. For example, *Sap*^-/-^ T cells, although capable of differentiating into early Tfh cells, are unable to be efficiently recruited to and retained in a nascent germinal center resulting an impaired germinal center reaction^14^. CXCR5 down-regulation and CXCR4 up-regulation result in re-distribution of Tfh cells from light zone to dark zone in GC, partially explaining the defects of GC responses and poor vaccine-induced immunity in older individuals^15^. As for Tfh cells, CXCR5 is also essential for B cells located in light zone of GC^16^, while CXCR4 is required for B cells migrating into dark zone of GC^17^. Although Bcl6 is the master transcription factor for Tfh cell differentiation, it does not directly regulate CXCR5 expression during Tfh cell differentiation especially during their migration into GC^18, 19^. Ascl2 initiates Tfh cell differentiation by directly up-regulating *Cxcr5* but not *Bcl6*^20^. However, how Tfh cells maintain and further up-regulate CXCR5 expression during pre-Tfh to GC-Tfh is still unknown.

The transcription factor Bhlhe40 (basic helix–loop–helix [bHLH] family member e40), also known as Bhlhb2, Dec1, or Stra13, is induced by TCR signaling^21^. Bhlhe40 has been reported to be critical for inducing autoimmune diseases, such as experimental autoimmune encephalomyelitis by regulating GM-CSF and IL-10^22^. Through inhibiting IL-10 production, Bhlhe40 helps balance inflammatory and anti-inflammatory type 1 immune responses^23, 24^. Bhlhe40 also regulates the differentiation and function of Th2^25^ and CD8 T cells^26^. It has been recently reported that Bhlhe40 restrains GC reaction by limiting activated CD4 T cells proliferation and the earliest generation of GC B cells^27^. However, the mechanism through which Bhlhe40 regulates Tfh cell differentiation is still elusive.

Pou2af1 (POU domain class 2-associating factor 1), also well known as Bob1, OBF-1 or OCA-B, has long been considered as a B cell-specific factor that interacts with the octamer transcription factors Oct1 and Oct2 to enhance octamer-dependent transcription^28^. Pou2af1 has also been reported to be involved in GC formation^29, 30, 31^ and Tfh function^32^. Since Pou2af1 is highly expressed in B cells and B cells are essential for Tfh cell differentiation, whether Pou2af1 regulates Tfh by regulating B cells, or whether there are T cell intrinsic effects of Pou2af1 on Tfh differentiation remains uncertain.

In this study, we found that Bhlhe40 suppressed Pou2af1 expression during T cell activation towards all Th cell fates. In the absence of Bhlhe40, Tfh cell differentiation was enhanced in a manner dependent on the upregulation of Pou2af1. Pou2af1 was critical for Tfh cell differentiation and migration into GC through upregulation of CXCR5 expression in a T cell intrinsic manner. On the other hand, Pou2af1 deficiency did not affect initial Bcl6 induction during early Tfh cell differentiation. Thus, our work highlights the importance of the Bhlhe40-Pou2af1 axis downstream of Bcl6-Blimp1 in regulating CXCR5 expression during cell maturation from pre-Tfh to GC-Tfh cells.

## RESULTS

### Bhlhe40 represses Pou2af1 during T cell differentiation

By re-analyzing our previously published RNA-Seq results^33^, we found that *Bhlhe40* expression was quickly induced under all four T helper (Th1, Th2, Th17 and iTreg) in vitro polarization conditions (Fig. 1A), similar to *Irf4* expression (fig. S1A). To assess whether Bhlhe40 is up-regulated at the protein level after TCR stimulation, we generated a V5-tagged Bhlhe40 mouse strain (fig. S1B), in which the DNA sequence encoding the V5 tag (GKPIPNPLLGLDST) was inserted into the exon5 of Bhlhe40 immediately before the stop codon. Naïve CD4 T cells from these mice were then differentiated under Th1 or Th2 conditions in vitro as previously described^34^. Anti-V5 staining indicated that Bhlhe40 expression at the protein level was also quickly induced by 6 hours after anti-CD3 and anti-CD28 stimulation under both Th1 and Th2 conditions (Fig. 1B, and fig. S1C). Due to the difficulty in generating Tfh cells in vitro, we measured Bhlhe40 expression in Tfh cells in vivo by immunizing wild type (WT) and V5-tagged mice with AS15/CFA for 2 weeks. Flow cytometry analyses of splenocytes from the immunized mice showed that among the antigen-specific CD4 T (CD19^-^CD4^+^CD44^hi^AS15^+^) cells, Bhlhe40 was highly expressed in non-Tfh (PD-1^-^CXCR5^-^) cells but not in Tfh (PD-1^+^CXCR5^+^) cells (Fig. 1C). A similar expression pattern of Bhlhe40 was noted in CD4 T cells harvested from the draining lymph nodes (fig. S1D). To identify genes that are regulated by Bhlhe40 in distinct subclasses of Th cells, we compared RNA-Seq analyses of *Bhlhe40*^fl/fl^ cells with *Bhlhe40*^fl/fl^-CD4Cre cells differentiated under four culture conditions (Th1, Th2, Th17 and iTreg). Fifty-five genes were up-regulated in the absence of Bhlhe40 in all four T helper subsets (Fig. 1D). Among these genes, six encode transcription factors with *Pou2af1* ranked at the top. Pou2af1 was reported to be essential for germinal center formation^29, 30, 31^ and may be required for Tfh cell differentiation and function^32^. Indeed, through RNA-Seq analyses of Tfh cells generated in vivo, Pou2af1 was found highly expressed in the PD-1^+^CXCR5^+^ population with the expression of many other Tfh signature genes (cluster 2) but not non-Tfh signature genes (cluster 1, Fig. 1E). To test whether Bhlhe40 can directly bind to the *Pou2af1* gene, we performed ChIP-Seq analysis of Bhlhe40 DNA binding with anti-V5 immunoprecipitation, using in vitro polarized Th cells. We found that Bhlhe40 indeed bound to the promoter of *Pou2af1* in all four Th subsets (Fig. 1F). Taken together, our results indicate that Bhlhe40 may repress *Pou2af1* expression in T helper subsets through a direct binding to its promoter. Given the opposite expression pattern of Bhlhe40 and Pou2af1 in Tfh and non-Tfh, we hypothesized that the Bhlhe40-Pou2af1 axis may play an important role during Tfh cell differentiation.

### Pou2af1 is required for Tfh cell differentiation

To address the function of Pou2af1 during Tfh cell differentiation, we generated a *Pou2af1* knockout mouse strain (*Pou2af1*^-/-^) by deleting exon3, exon4, and most of exon2 and exon5 as previously described^29^ via CRISPR-Cas9 technology (fig. S2A). Consistent with the previous reports^30, 31^, B cells but not T cells in *Pou2af1*^-/-^ mice were found reduced as expected (Fig. 2A and fig. S2B). We then immunized WT and *Pou2af1*^-/-^ mice with AS15/CFA for 2 weeks, and found that Tfh (PD-1^+^CXCR5^+^) cells were reduced in draining lymph nodes of *Pou2af1*^-/-^ mice (Fig. 2, B and C). Since B cells are known to be involved in Tfh cell generation, to exclude the effect of B cell defect in *Pou2af1*^-/-^ mice, we constructed bone marrow (BM) chimeric animals in which the vast majority of B cells were normal but T cells were from either WT or *Pou2af1*^-/-^ (Fig. 2D). Six weeks after BM reconstitution, we confirmed that B cells in different groups were indeed identical (Fig. 2E, and fig. S2C). However, Tfh cells were still reduced in *Pou2af1*^-/-^ chimeras compared with WT mixed chimeras (Fig. 2, F and G) demonstrating a T cell intrinsic defect in *Pou2af1*^-/-^ cells to become Tfh cells.

### Bhlhe40 represses Tfh cell differentiation

Given that Bhlhe40 represses Pou2af1 and that Pou2af1 is required for Tfh cell differentiation, Bhlhe40 could be involved in repressing Tfh cell formation. Indeed, one week after immunization with AS15/CFA, we found that AS15-specific Tfh (PD-1^+^CXCR5^+^ or Bcl6^+^CXCR5^+^) cell were increased in both draining lymph nodes (Fig. 3, A and B) and spleen (fig. S3A) from the Bhlhe40 knockout mice. To exclude Bhlhe40 function in other types of cells, we performed similar experiments with *Bhlhe40*^fl/fl^-CD4Cre mice, which lack Bhlhe40 expression in T cells. Tfh cells were also increased in *Bhlhe40*^fl/fl^-CD4Cre mice compared with WT mice 5 days post immunization (Fig. 3, C and D). However, after 2 weeks, the difference in Tfh cells between immunized WT and *Bhlhe40*^fl/fl^-CD4Cre became smaller (fig. S3, B and C), which may be explained by the fact that Bhlhe40 is down-regulated during Tfh cell differentiation in WT mice.

### Mutually exclusive expression of Bhlhe40 and CXCR5 during T cell response in vivo

To further assess the expression levels of *Bhlhe40* during Tfh cell differentiation in vivo, we immunized Bhlhe40-V5 mice and then stained CD4 T cells from the spleen using anti-V5, which reflects Bhlhe40 expression. Within the antigen-specific CD4 T (B220^-^CD4^+^CD44^hi^AS15^+^) cells, Bhlhe40 and CXCR5 showed a mutually exclusive expression pattern, i.e., CXCR5^+^ cells were V5^-^ but V5^+^ cells were CXCR5^-^ (Fig. 4A). On the other hand, Bhlhe40 and Bcl6 can be co-expressed since a small Bcl6^+^V5^+^ population was detected. While Bhlhe40 expression was detected in non-Tfh (Bcl6^-^CXCR5^-^) cells, Tfh cells (Bcl6^+^CXCR5^+^) did not express Bhlhe40 (Fig. 4B). Furthermore, Bhlhe40-V5 expression was also found in Bcl6^+^CXCR5^-^ population (Fig. 4B). These results indicate that Bhlhe40 may play an important role in limiting CXCR5 expression independent of the Bcl6-Blimp1 axis.

To facilitate studying the expression and functions of Pou2af1, we generated Pou2af1-HA-tdTomato reporter mouse strain through CRISPR-Cas9. In these mice, tdTomato can reflect Pou2af1 expression, whereas the HA-tag on N-terminal of Pou2af1 can be used to pull down Pou2af1 (fig. S4A). Using these reporter mice, we found that Pou2af1 was highly expressed on a subset of B cells as expected (fig. S4B), which is also consistent with a previous report^28^. A portion of B cells from Pou2af1-HA-tdTomato reporter mice (Pou2af1-HA-TG) expressed tdTomato, and the proportion was higher in CXCR5^+^ B cells (36%) than what was in CXCR5^-^ B cells (8.3%) (fig. S4B). We also bred these reporter mice to *Bhlhe40*^fl/fl^-CD4Cre mice to generate Pou2af1-tdTomato-*Bhlhe40*^fl/fl^-CD4Cre mice. Furthermore, expression of Pou2af1-tdTomato was increased after knocking out Bhlhe40 in cells cultured under Th1 conditions (fig. S4, C and D).

### Pou2af1- and Bhlhe40-mediated gene regulation during Tfh cell differentiation

To gain more insights on how Bhlhe40 and Pou2af1 regulate Tfh cell differentiation, we performed RNA-Seq as shown in fig. S5A. By adoptively transferring bone marrow from WT, *Pou2af1*^-/-^, or *Bhlhe40*^-/-^ mice together with that from *TCRα*^-/-^ mice with 1:5 ratio and then immunizing the mice, we were able to study the T cell intrinsic effect of Pou2af1 and Bhlhe40 during Tfh cell differentiation 2 weeks post-immunization. Three sub-populations (PD-1^-^ CXCR5^-^, PD-1^-^CXCR5^+^, PD-1^+^CXCR5^+^) among the antigen-specific CD4 T cells were sorted and subjected to RNA-Seq analyses (fig. S5B). Initial analysis of the RNA-Seq results confirmed expected genetic deletions in cells from different groups (fig. S5C).

As shown in Fig. 1E, we first identified 887 different expression genes (DEGs) within the WT groups (including Tfh signature genes and non-Tfh signature genes), which distinguish three populations of antigen specific effector CD4 T cells populations (PD-1^-^CXCR5^-^, PD-1^+^CXCR5^-^, PD-1^+^CXCR5^+^). Comparing WT and *Pou2af1*^-/-^ PD-1^+^CXCR5^+^ groups, 186 DEGs were identified, among which 173 of them overlapped with the 887 signature genes (Fig. 5A). CXCR5 was found reduced in *Pou2af1*^-/-^ compared with WT (Fig. 5B). Strikingly, all 126 down-regulated genes in the absence of Pou2af1 were Tfh signature genes whereas all 47 up-regulated genes were non-Tfh-signature genes (Fig. 5C). Furthermore, we found increased Pou2af1 expression in the PD-1^+^CXCR5^-^ population in the absence of Bhlhe40 (Fig. 5D), suggesting that downregulation of Bhlhe40 and thus the induction of Pou2af1 expression allows GC Tfh generation through upregulating CXCR5 (Fig. 5E). To test whether Pou2af1 can directly regulate CXCR5, we performed anti-HA ChIP-Seq analysis of B cells harvested from Pou2af1-HA-tdTmato reporter mice. A Pou2af1 binding peak 5’ of the *Cxcr5* gene that contains an Oct1/2 motif (ATGCAAA) was identified (Fig. 5F).

### Pou2af1 regulates Tfh cell differentiation downstream of Bhlhe40

To confirm the relationship between Pou2af1 and Bhlhe40 in regulating Tfh cell development, we crossed *Pou2af1*^-/-^ mice with *Bhlhe40*^-/-^ mice to generate *Pou2af1*^-/-^*Bhlhe40*^-/-^ mice. By comparing immunized WT, *Pou2af1*^-/-^, and *Pou2af1*^-/-^*Bhlhe40*^-/-^ mice, we found that Tfh cells (CXCR5^+^PD-1^+^) were similarly reduced in *Pou2af1*^-/-^*Bhlhe40*^-/-^ mice as in *Pou2af1*^-/-^ mice (Fig. 6, A and B). Since B cells in *Pou2af1*^-/-^*Bhlhe40*^-/-^ mice were also found reduced, we performed similar BM Chimeras experiments as shown in Fig. 2E to exclude the effect of B cell defect. Tfh cells (CXCR5^+^Bcl6^+^) were still reduced in *Pou2af1*^-/-^*Bhlhe40*^-/-^ group (Fig. 6, C and D) while B cells in these WT, *Pou2af1*^-/-^ and *Pou2af1*^-/-^*Bhlhe40*^-/-^ chimeras were similar (fig. S6). Consistent with the idea that Bhlhe40-Pou2af1 axis mainly regulates CXCR5 expression independent of Bcl6, the Bcl6 ^+^ population remained similar among WT, *Pou2af1*^-/-^ and *Pou2af1*^-/-^*Bhlhe40*^-/-^ group with accumulation of CXCR5^-^Bcl6^+^ population when both Bhlhe40 and Pou2af1 were absent (Fig. 6, C and D). Since *Pou2af1*^-/-^*Bhlhe40*^-/-^ largely showed a similar phenotype as *Pou2af1*^-/-^, we conclude that Pou2af1 functions downstream of Bhlhe40 in regulating Tfh cell differentiation.

### Optimal CXCR5 expression is required for Tfh cell migration into germinal center

CXCR5 is well-known to be involved in Tfh migration from T-B border to GC. Our data have shown that the Bhlhe40-Pou2af1 axis play an important role in regulating CXCR5 expression downstream of Bcl6. To test whether such optimal regulation of CXCR5 expression by Bhlhe40 and Pou2af1 has any impact on cell migration in a cell intrinsic manner, we designed another mixed bone marrow experiment in which a mixture of WT BM and BM from different genetic background was adoptive transferred into irradiated *Rag1*^-/-^ mice (Fig. 7A). Seven weeks after BM reconstitution, these *Rag1*^-/-^ recipients were immunized with AS15/CFA. One week later, Tfh cells from the draining lymph nodes were analyzed. As expected, in ‘WT (CD45.2^+^) +WT (CD45.2^-^)’ group, the MFI of CXCR5 expression by both WT AS15^+^ CD4 T cells was similar (Fig. 7B). However, in ‘*Pou2af1*^-/-^ (CD45.2^+^) +WT (CD45.2^-^)’ group, the MFI of CXCR5 was reduced in *Pou2af1*^-/-^ population compared with WT AS15^+^ CD4 T cells in the same animal. In contrast, the MFI of CXCR5 was increased in *Bhlhe40*^-/-^ population compared with WT AS15^+^ CD4 T cells in ‘*Bhlhe40*^-/-^ (CD45.2^+^) +WT (CD45.2^-^)’ group. Furthermore, the MFI of CXCR5 was again decreased in *Pou2af1*^-/-^*Bhlhe40*^-/-^ population compared with the WT counterpart in ‘*Pou2af1*^-/-^*Bhlhe40*^-/-^ (CD45.2^+^) +WT (CD45.2^-^)’ group. There was a progressive reduction in the proportion of *Pou2af1*^-/-^ cells relative to the WT cells with increased CXCR5 expression (Fig. 7C). A similar reduction was observed in the *Pou2af1*^-/-^*Bhlhe40*^-/-^ group reaffirming that Pou2af1 functions downstream of Bhlhe40 in regulating proper CXCR5 expression in a T cell intrinsic manner.

**Fig. 1.**
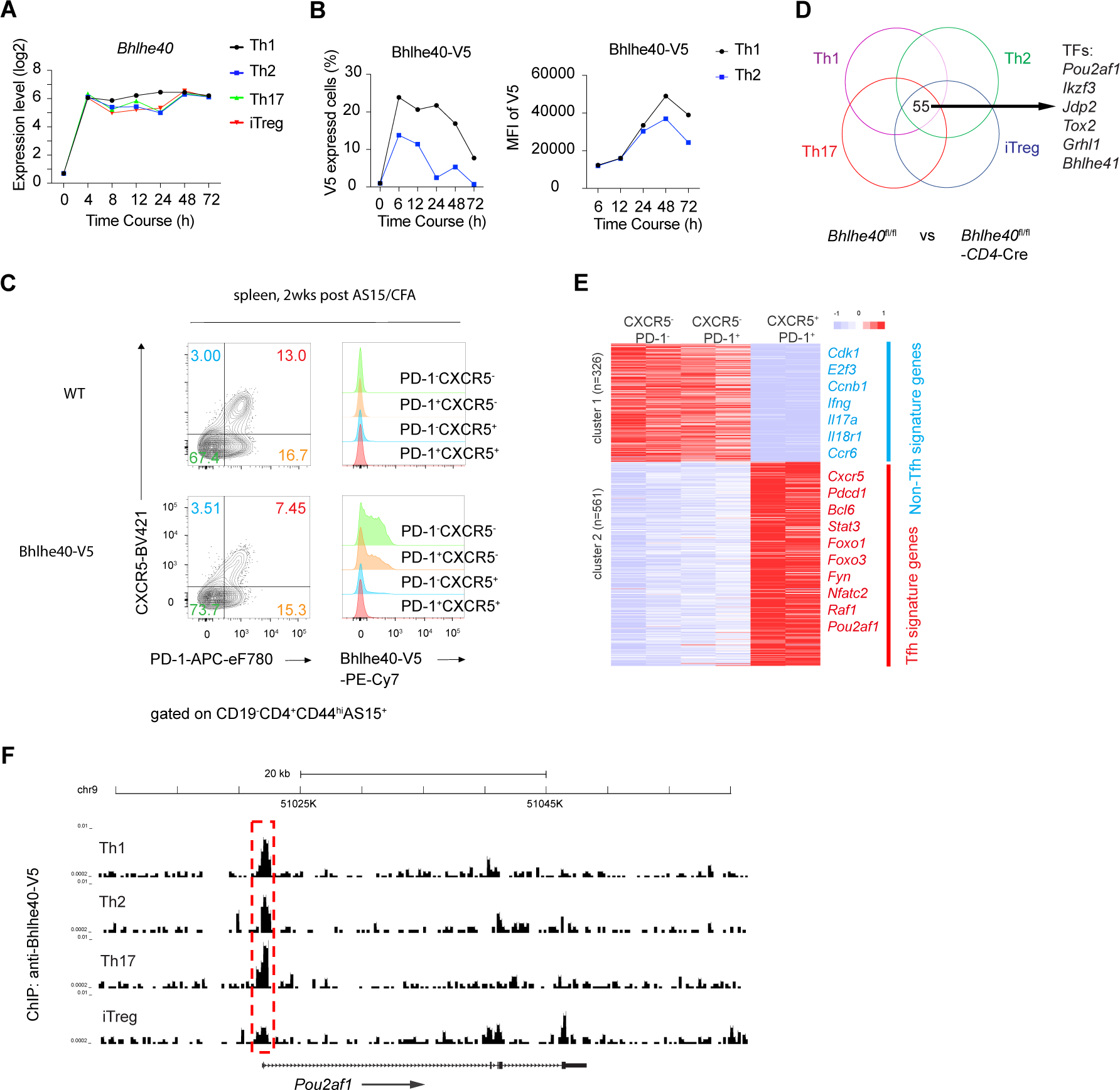
Bhlhe40 represses Pou2af1 in conventional T cells. (**A**) RNA-Seq data^33^ were re-analyzed for *Bhlhe40* expression during Th1, Th2, Th17 and iTreg culture in vitro. (**B**) Naïve CD4 T cells were purified from Bhlhe40-V5 tagged mice and primed under Th1 or Th2 conditions. Bhlhe40 expression was assessed by anti-V5 staining at indicated time points. (**C**) WT and Bhlhe40-V5 tagged mice were immunized (s.c.) with AS15/CFA for 2 weeks (2wks), and the expression of *Bhlhe40* by the splenic AS15-specific effective CD4 T cells in four quadrants (PD-1^-^CXCR5^-^, PD-1^+^CXCR5^-^, PD-1^-^CXCR5^+^, PD-1^+^CXCR5^+^) was assessed by flow cytometry. (**D**) Naïve CD4 T cells from *Bhlhe40*^fl/fl^ or *Bhlhe40*^fl/fl^-CD4Cre mice were cultured under Th1, Th2, Th17 and iTreg conditions, then harvested for RNA-Seq analysis. (**E**) AS15-specific wide type (WT) CD4 T cells (B220^-^CD4^+^CD44^hi^AS15^+^) were sorted for three populations (PD-1^-^CXCR5^-^, PD-1^+^CXCR5^-^, PD-1^+^CXCR5^+^) from inguinal lymph nodes and subjected to RNA-Seq analyses. Samples were in biological duplicates. (**F**) Naïve CD4 T cells from Bhlhe40-V5 tagged mice were cultured under Th1, Th2, Th17 and iTreg conditions, then harvested for ChIP-Seq analysis using anti-V5. Samples (D to F) were in biological duplicates. Data are representative of two (B and C) independent experiments.

**Fig. 2.**
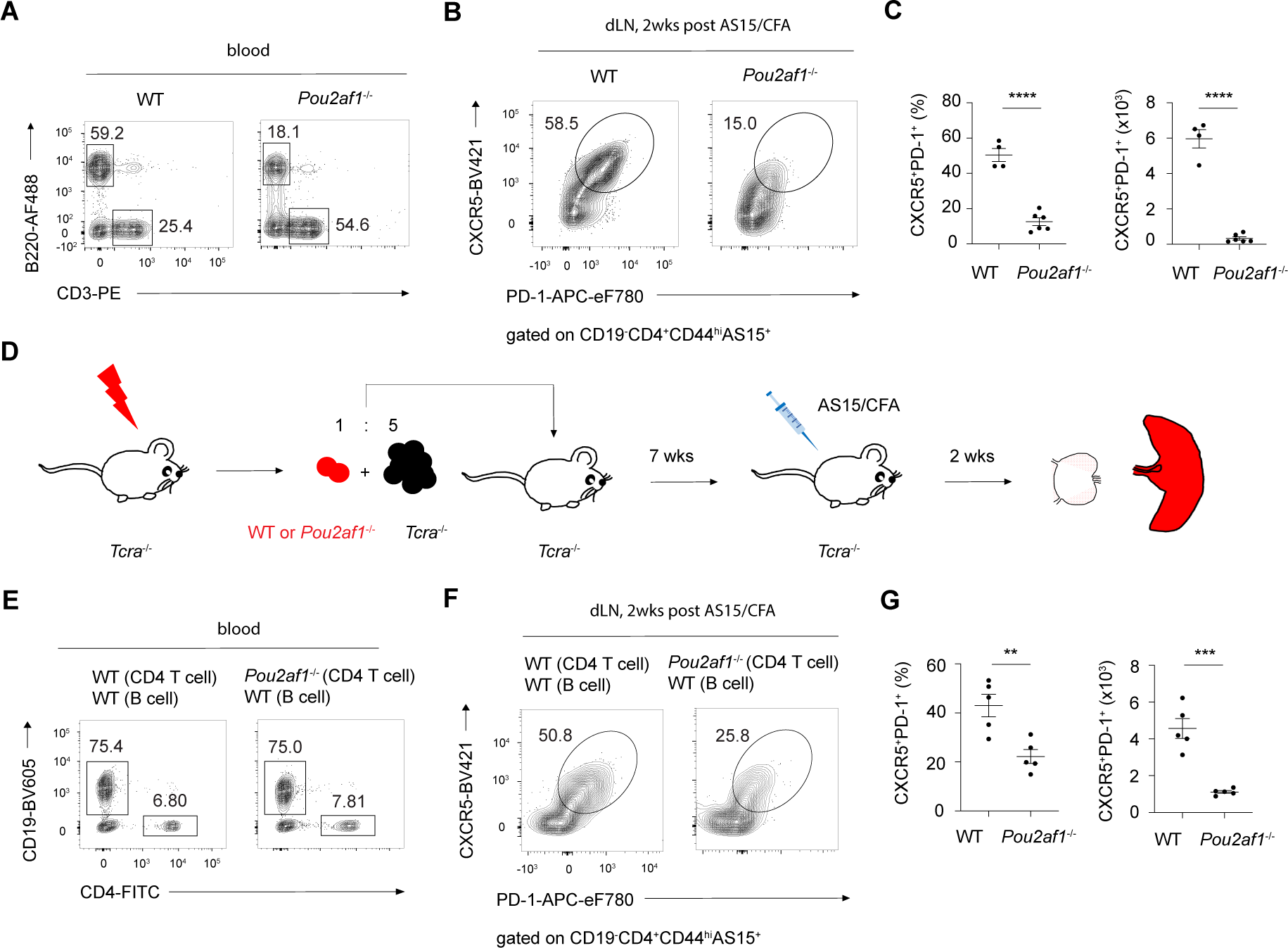
Pou2af1 is required for Tfh cell differentiation. (**A**) Blood cells from WT or *Pou2af1*^-/-^ mice were stained for T cells (B220^-^CD3^+^) and B cells (B220^+^CD3^-^) by flow cytometry. (**B**) WT and *Pou2af1*^-/-^ mice were immunized (s.c.) with AS15/CFA for 2 weeks, and AS15-specific CD4 T cells (CD19^-^CD4^+^CD44^hi^AS15^+^) from inguinal lymph nodes were analyzed for Tfh (PD-1^+^CXCR5^+^) cells. (**C**) Summary of differences in percentage and cell numbers of Tfh cells between WT (*n*=4) and *Pou2af1*^-/-^ (*n*=6) in (B). (**D**) Experimental procedure of immunizing bone marrow chimeras *TCRα*^-/-^ mice adoptively transferred a bone marrow mixture of *TCRα*^-/-^ and WT or *Pou2af1*^-/-^ mice with AS15/CFA. (**E**) Blood cells from (D) were assessed for CD4 T cells (CD19^-^CD4^+^) and B cells (CD19^+^CD4^-^) by flow cytometry. (**F)** Tfh cells in inguinal lymph nodes from (D) were analyzed as (B). (**G**) Summary of percentage and cell number of Tfh cells from WT group (*n*=5) and *Pou2af1*^-/-^ group (*n*=5) in (D). * p < 0.05, ** p < 0.01, *** p < 0.001, **** p < 0.0001. Error bars indicate SEM. Data are representative of more than three (A-C) and two (D-G) independent experiments.

**Fig. 3.**
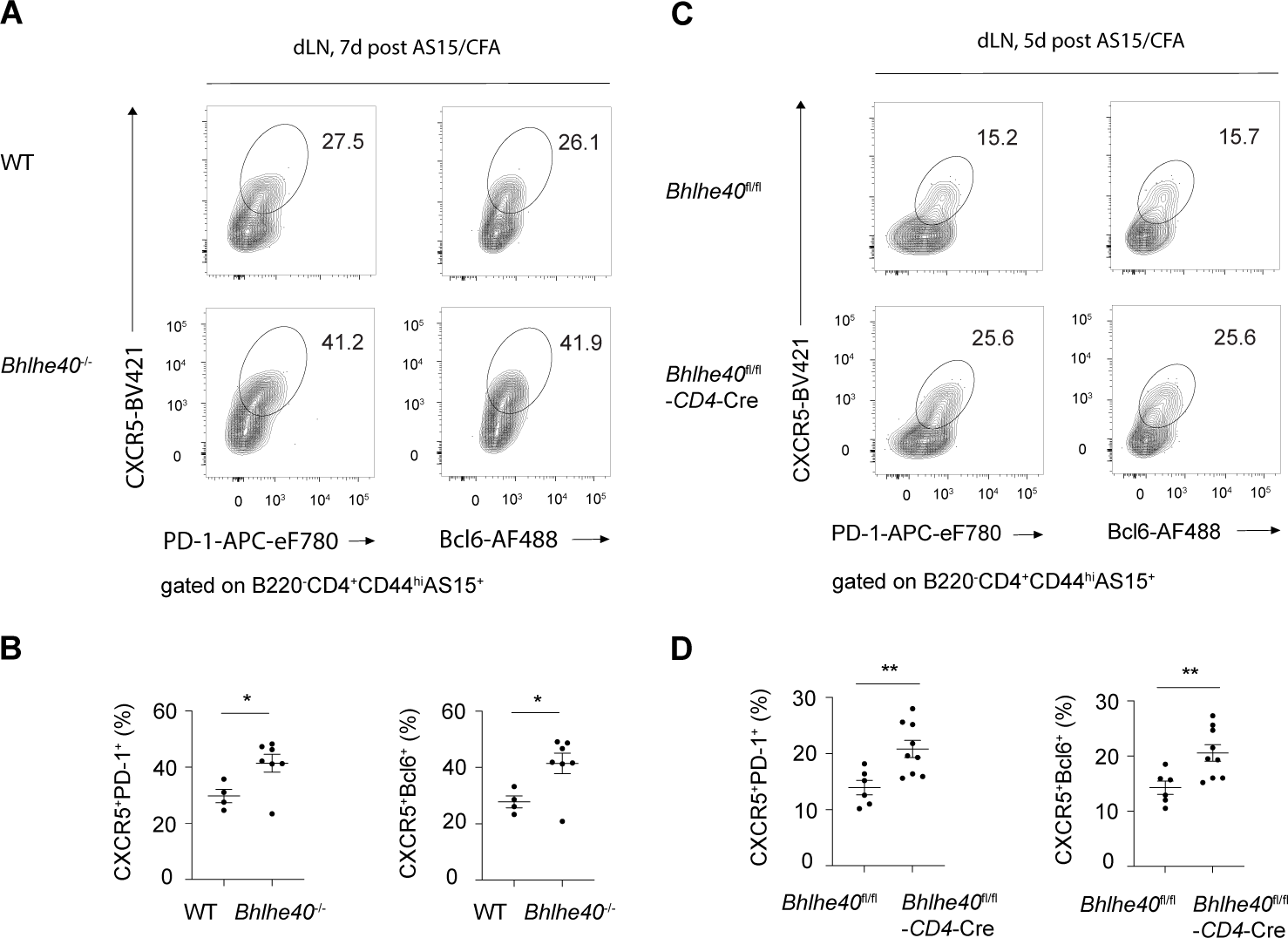
Bhlhe40 represses Tfh cell differentiation. (**A**) WT (*n*=4) and *Bhlhe40*^-/-^ (*n*=7) mice were immunized (s.c.) with AS15/CFA for 7d, and AS15-specific CD4 T cells from inguinal lymph nodes were analyzed for Tfh (PD-1^+^CXCR5^+^ or Bcl6^+^CXCR5^+^) cells by flow cytometry. (**B**) Summary of percentage difference in Tfh cells between WT and *Bhlhe40*^-/-^ in (A). (**C**) WT (*n*=6) and *Bhlhe40*^fl/fl^-CD4Cre (*n*=9) mice were immunized (s.c.) with AS15/CFA for 5d, and AS15-specific effective CD4 T cells from inguinal lymph nodes were analyzed for Tfh (PD-1^+^CXCR5^+^ or Bcl6^+^CXCR5^+^) cells by flow cytometry. (**D)** Summary of percentage difference in Tfh cells between WT and *Bhlhe40*^fl/fl^-CD4Cre in (C). * p < 0.05, ** p < 0.01, *** p < 0.001, **** p < 0.0001. Error bars indicate SEM. Data are representative of two (A-D) independent experiments.

**Fig. 4.**
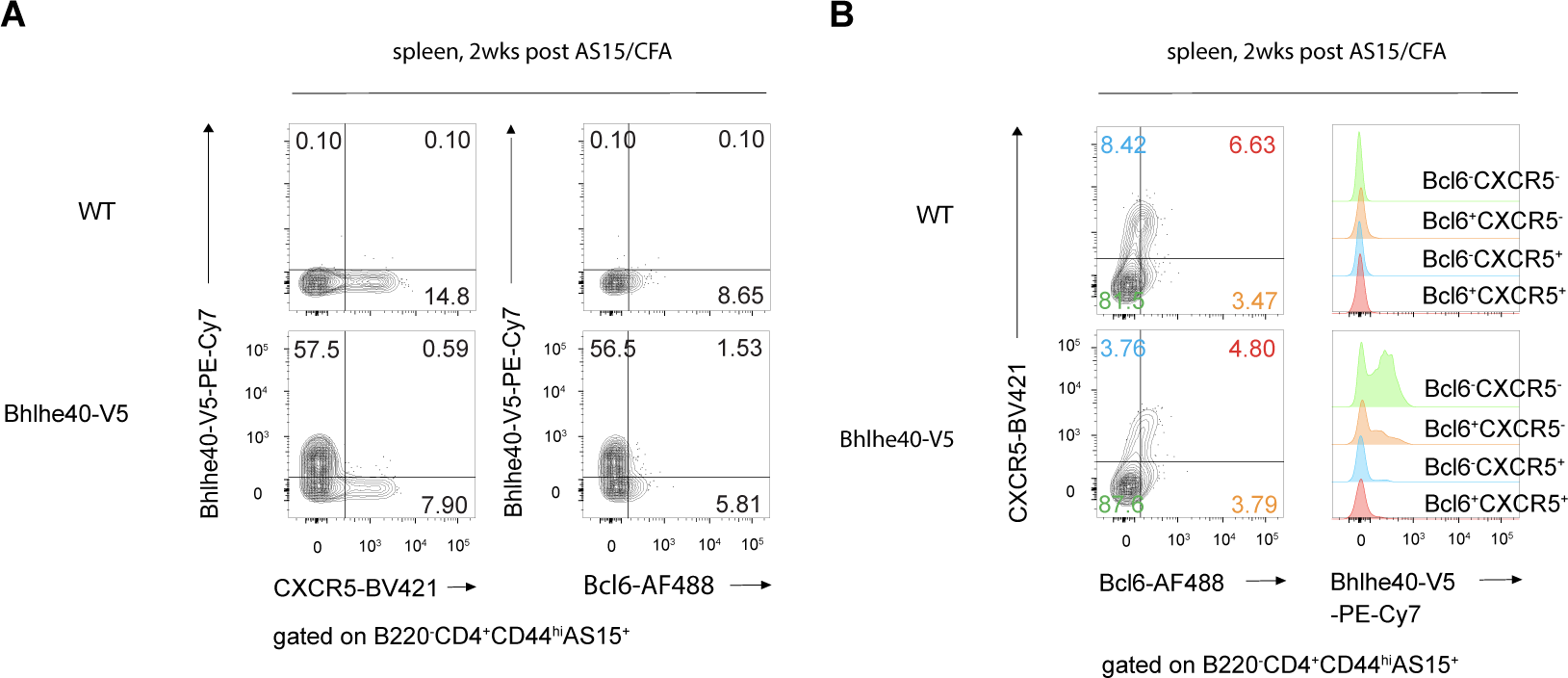
Bhlhe40 and CXCR5 expression is mutual exclusive. (**A** and **B**) WT and Bhlhe40-V5 mice were immunized (s.c.) with AS15/CFA for 2 weeks, the splenic AS15-specific CD4 T cells were assessed by flow cytometry. (A) Flow cytometry plots showing the expression pattern between CXCR5 and Bhlhe40-V5, or Bcl6 and Bhlhe40-V5. (B) The expression of Bhlhe40 in four subpopulations (Bcl6^-^CXCR5^-^, Bcl6^+^CXCR5^-^, Bcl6^-^CXCR5^+^, Bcl6^+^CXCR5^+^) was assessed as indicated. Data are representative of two (A-B) independent experiments.

**Fig. 5.**
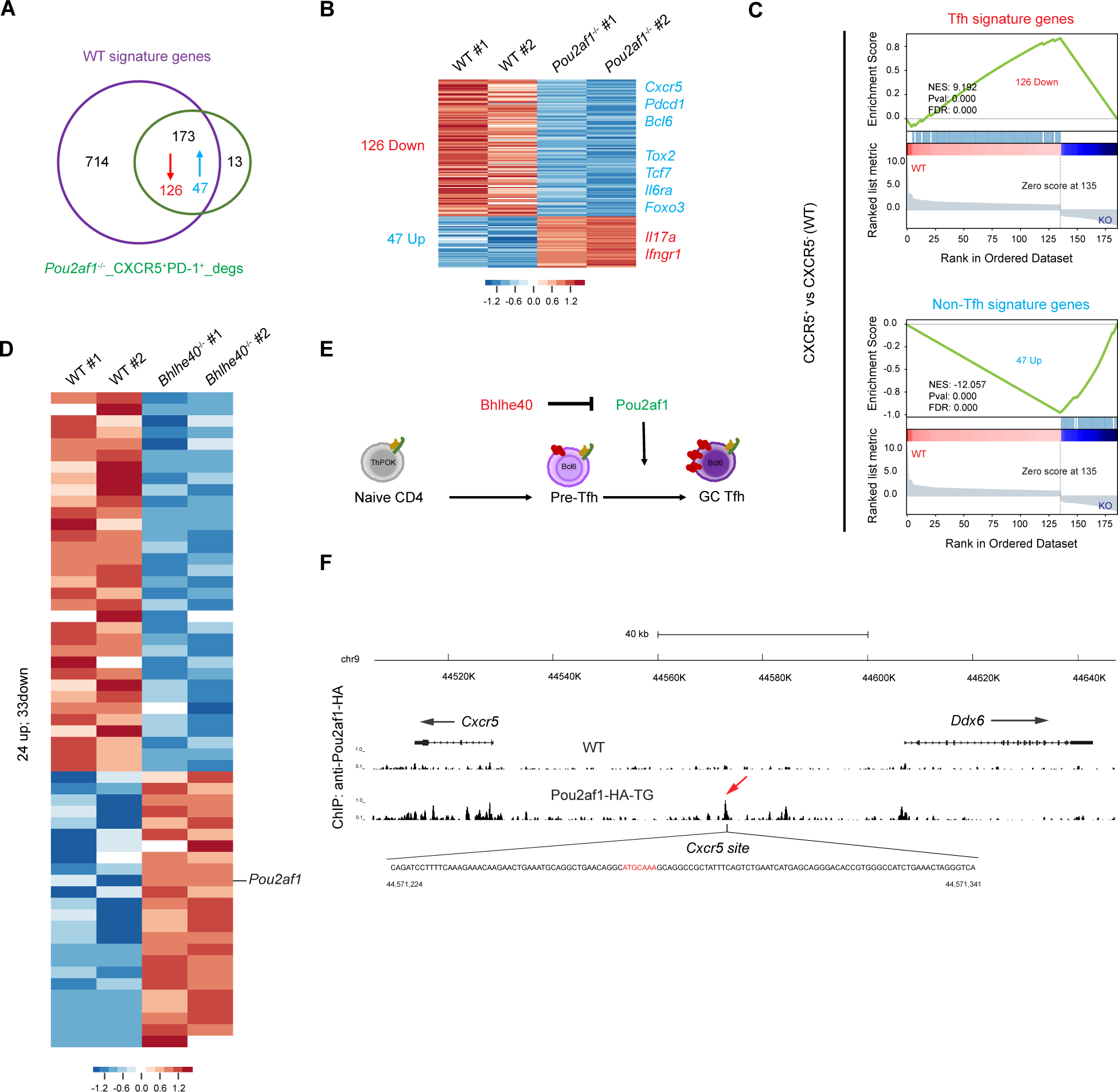
Pou2af1 and Bhlhe40 regulate Tfh cell differentiation in cell intrinsic manner. (**A** to **D**) AS15-specific WT, *Pou2af1*^-/-^, *Bhlhe40*^-/-^ CD4 T cells (B220^-^CD4^+^CD44^hi^AS15^+^) were sorted for three populations (PD-1^-^CXCR5^-^, PD-1^+^CXCR5^-^, PD-1^+^CXCR5^+^) from inguinal lymph nodes and used for RNA-Seq analysis. Samples are in biological duplicates. Experimental procedure was shown in SM Fig. 5A. (A) Overlap between signature genes shown in Fig. 1E and DEGs (different expression genes) between WT and *Pou2af1*^-/-^ CXCR5^+^PD-1^+^ cells. (B) Heat map of 126 down-regulated genes and 47 up-regulated genes among the DEGs in (A). (C) GSEA of the 126 down-regulated genes and 47 up-regulated genes in (B). (D) DEGs between WT and *Bhlhe40*^-/-^ CXCR5^-^PD-1^+^ cells. (**E**) A model for Bhlhe40- and Pou2af1-mediated regulation of Tfh cell differentiation. (**F**) Splenic B cells were sorted from WT or Pou2af1-HA-TG mice, and anti-HA was used for ChIP-Seq analysis for Pou2af1 binding in the genome. An Oct1/2 binding octamer DNA motif^28, 54^ (red) was found in *Cxcr5* site.

**Fig. 6.**
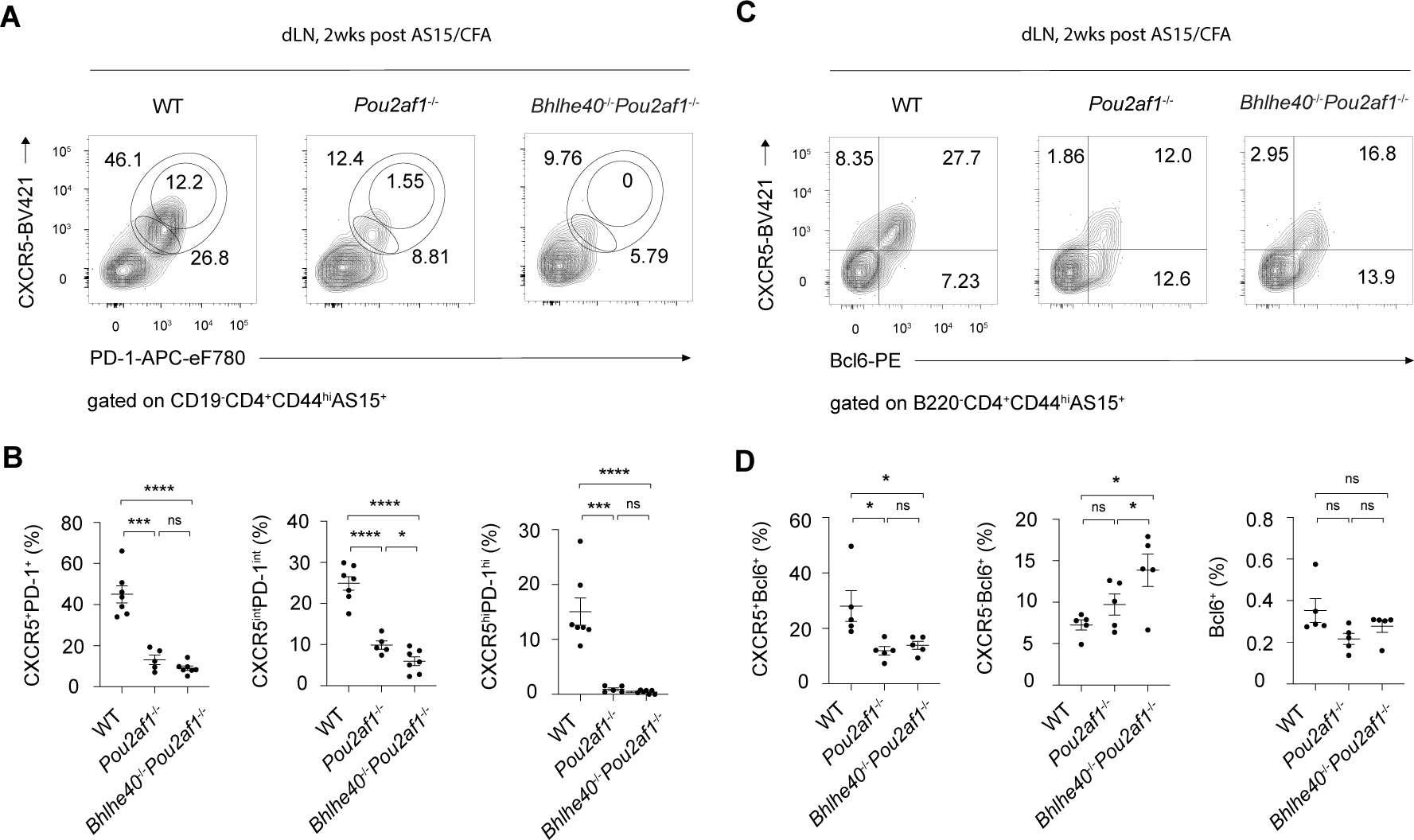
Pou2af1 functions downstream of Bhlhe40 during Tfh cell differentiation. (**A**) WT (*n*=7), *Pou2af1*^-/-^ (*n*=5), *Bhlhe40*^-/-^*Pou2af1*^-/-^ (*n*=7) mice were immunized (s.c.) with AS15/CFA for 2 weeks, and AS15-specific CD4 T cells from inguinal lymph nodes were analyzed for PD-1^int^CXCR5^int^, PD-1^hi^CXCR5^hi^ and PD-1^+^CXCR5^+^ (including both PD-1^int^CXCR5^int^ and PD-1^hi^CXCR5^hi^) by flow cytometry. (**B**) Summary of percentage of PD-1^+^CXCR5^+^, PD-1^int^CXCR5^int^, PD-1^hi^CXCR5^hi^ cells in (A). (**C**) AS15-specific CD4 T cells from inguinal lymph nodes of WT (*n*=5), *Pou2af1*^-/-^ (*n*=5), *Bhlhe40*^-/-^*Pou2af1*^-/-^ (*n*=5) group in bone marrow chimeras model as Fig. 2E were analyzed for Bcl6^+^CXCR5^+^, Bcl6^+^CXCR5^-^ and Bcl6^+^ (including both Bcl6^+^CXCR5^+^ and Bcl6^+^CXCR5^-^) by flow cytometry. (**D**) Summary of percentage of Bcl6^+^CXCR5^+^, Bcl6^+^CXCR5^-^ and Bcl6^+^ cells in (C). * p < 0.05, ** p < 0.01, *** p < 0.001, **** p < 0.0001. Error bars indicate SEM. Data are representative of three (A and B) and two (C and D) independent experiments.

**Fig. 7.**
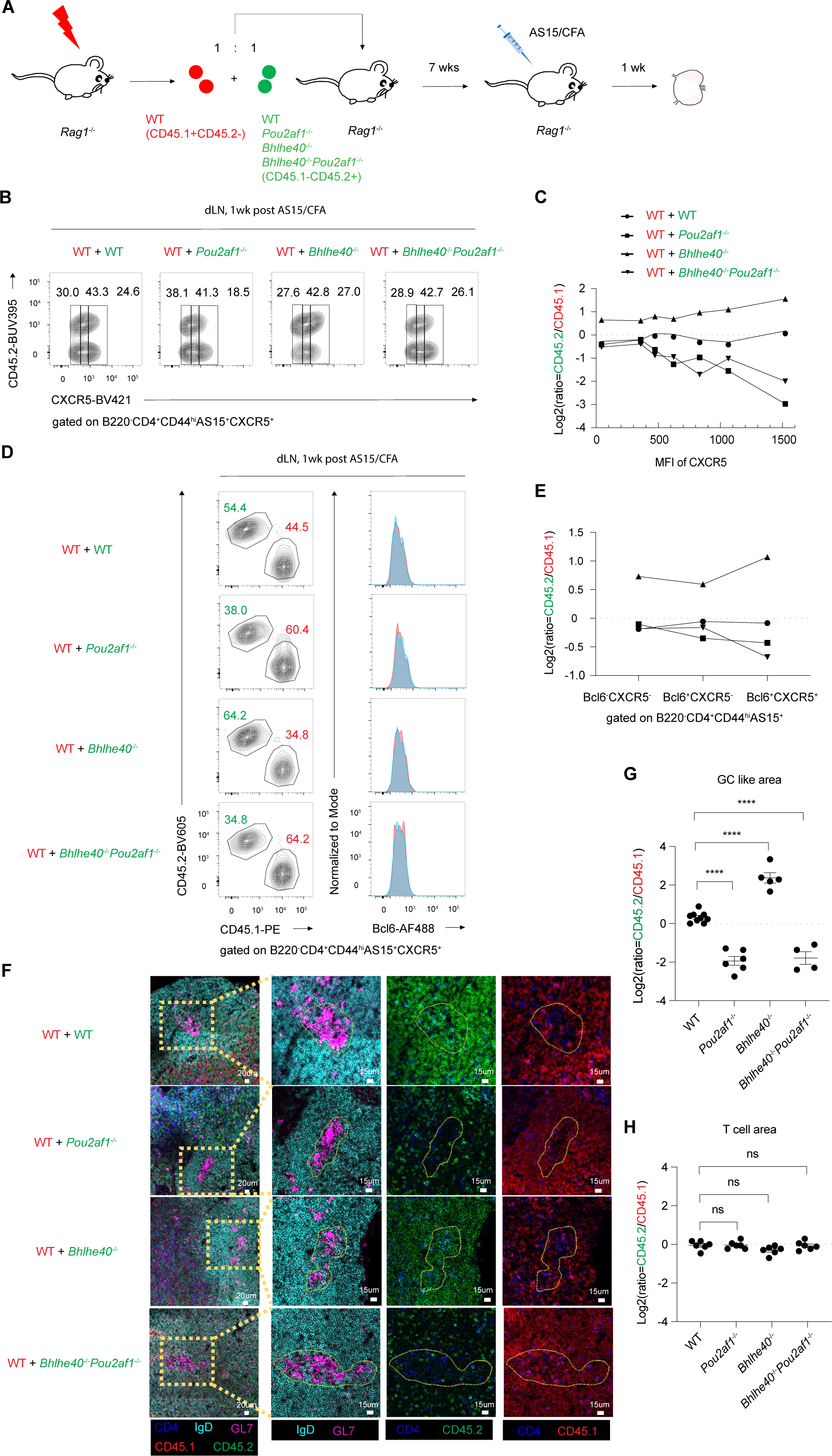
Regulation of optimal CXCR5 level through Bhlhe40-Pou2af1 axis during Tfh cell maturation. (**A**) Experimental procedure of immunizing mixed bone marrow *Rag1*^-/-^ mice by adoptive transferring bone marrow mixture of WT (CD45.1^+^CD45.2^-^) (red) mice with indicated genotyped (WT, *n*=3; *Pou2af1*^-/-^, n=*2*; *Bhlhe40*^-/-^, n=*3*; *Bhlhe40*^-/-^*Pou2af1*^-/-^, *n*=3) mice (CD45.1^-^ CD45.2^+^) (green) mice with AS15/CFA for one week. (**B**) B220^-^CD4^+^CD44^hi^AS15^+^CXCR5^+^ cells from inguinal lymph nodes were assessed by flow cytometry in four different groups from (A). (**C**) Summary of the cell ratios (indicated genotype/WT in the same mouse) based on different CXCR5 MFIs. (**D**) Same cells shown in (B) were assessed for Bcl6 expression by flow cytometry. (**E**) Summary of the cell ratios (indicated genotype/WT in the same mouse) among three cell populations (Bcl6^-^CXCR5^-^, Bcl6^+^CXCR5^-^, Bcl6^+^CXCR5^+^). (**F**) Inguinal lymph nodes from (A) were fixed and stained for confocal image. Images in yellow box of left were further enlarged to show more details in germinal center area (GL7^+^). Anti-CD4 (blue) and anti-IgD (cyan) were used to show T cell area and B cell area. Anti-GL7 (purple) was used to show germinal center. Anti-CD45.2 was used to show different indicated genotyped adoptive cells, and anti-CD45.1 (red) was used to show WT adoptive cells from the same mouse. (**G**) Summary of ratio of indicated genotyped Tfh cells (CD4^+^CD45.2^+^)/WT Tfh cells (CD4^+^CD45.1^+^) in GC like area in (F) random GC like areas were chosen from four indicated genotyped inguinal lymph nodes slice samples (WT, *n*=9; *Pou2af1*^-/-^, *n*=6; *Bhlhe40*^-/-^, *n*=5; *Bhlhe40*^-/-^*Pou2af1*^-/-^, *n*=4). (**H**) Same analysis as in (G) but for T cell area. Random T cell areas were chosen from four indicated genotyped inguinal lymph nodes slice samples (WT, *n*=6; *Pou2af1*^-/-^, *n*=6; *Bhlhe40*^-/-^, *n*=6; *Bhlhe40*^-/-^*Pou2af1*^-/-^, *n*=6). * p < 0.05, ** p < 0.01, *** p < 0.001, **** p < 0.0001. Error bars indicate SEM. Data are representative of two (A to H) independent experiments.

To test whether Bhlhe40 or Pou2af1 deficiency affects Bcl6 expression in this chimeric model, we further analyzed Bcl6 expression in antigen specific CD4 T cells from different groups compared with their WT counterparts in the same mice. Consistent with the results shown in Fig. 6D, the MFI of Bcl6 expression was similar between WT and gene deficient cells in the same animal (Fig. 7D). Thus, Bhlhe40-Pou2af1 regulates CXCR5 independent of Bcl6 induction. Consequently, an obvious change in ratio between CD45.1 and CD45.2 population was observed only when cells progressed from Bcl6^+^CXCR5^-^ to Bcl6^+^CXCR5^+^ in the DKO chimeras (Fig. 7E).

To assess whether optimal expression of CXCR5 has any effect on cell migration to GC, we used these immunized mixed BM chimeric mice for confocal imaging. As shown in Fig. 7F and 7G, in ‘WT (CD45.2^+^) +WT (CD45.2^-^)’ group, we found equal numbers of (CD45.2^+^) and (CD45.2^-^) GC-Tfh cells. However, in ‘*Pou2af1*^-/-^ (CD45.2^+^) +WT (CD45.2^-^)’ group, most of the GC-Tfh cells were originated from WT (CD45.2^-^) group. By contrast, *Bhlhe40*^-/-^ (CD45.2^+^) cells were more potent than WT (CD45.2^-^) cells in forming GC-Tfh cells. Again, in ‘*Pou2af1*^-/-^ *Bhlhe40*^-/-^ (CD45.2^+^) +WT (CD45.2^-^)’ group, most of the GC-Tfh were from the WT (CD45.2^-^) group. While the ratio of CD45.2/CD45.1 within GC-Tfh population varied among these chimeras, CD45.2/CD45.1 ratio within CD4 T cells located in T cell area remained constant (Fig. 7H and fig. S7A), indicating that Bhlhe40 and Pou2af1 modulates Tfh cell migration into geminal center in a CD4 T cell intrinsic manner, which is likely due to effects on CXCR5 expression (fig. S7B).

## DISCUSSION

Tfh cells respond to chemokine CXCL13 through the expression of cell surface marker CXCR5, which is crucial for Tfh migration into B cell follicle and then germinal center to become GC-Tfh cells^35, 36, 37^. While the Bcl6-Blimp1 axis is critical for Tfh cell differentiation and lineage commitment, Bcl6-independent regulation of CXCR5 expression has been noted. During earlier Tfh cell differentiation primed by DCs, the transcription factor Ascl2 induces CXCR5 expression^20^. However, what regulates CXCR5 expression at a later stage of Tfh cell maturation is still unknown. Our results here demonstrate an optimal regulation of CXCR5 by the Bhlhe40-Pou2af1 axis during the pre-Tfh to GC-Tfh transition. Bhlhe40 negatively regulates CXCR5 via suppressing Pou2af1 expression. Such regulation is after Tfh cell lineage commitment which is determined by the Bcl6-Blimp1 axis.

Since Bhlhe40 has been reported to selectively restrict the generation of the earliest GC B cells^27^, and Pou2af1 is required for GC B cells generation^38, 39^, it is likely that the Bhlhe40-Pou2af1-medicated CXCR5 in B cells may also play an important role in the generation of GC B cells. T-bet can interact with Bhlhe40 in T cells^40^ and T-bet can also inhibit Tfh cell differentiation. However, whether T-bet-mediated suppression of Tfh cell differentiation involves Bhlhe40 is unknown.

Several pathways and molecules have been shown to regulate Bhlhe40 expression: TCR signaling can rapidly induce Bhlhe40 expression; Cytokines such as IL-1β can induce Bhlhe40 expression in T cells^41^; GATA3 may regulate Bhlhe40 expression in pathogenic Th17 cells of EAE model^42^. However, more work regarding Bhlhe40 regulation during Tfh cell differentiation is still needed. Similarly, while Bhlhe40 can suppress Pou2af1 expression, what signal(s) induces its expression is unclear. Furthermore, while we observed an important T cell intrinsic function of Pou2af1 in regulating CXCR5 expression, we can only detect a small fraction of Tfh cells expressing Pou2af1 at a given moment. It is likely that Pou2af1 expression in T cells is transient but its expression in B cells is more stable. Future fate-mapping experiments may clarify this point.

Bhlhe40 has other functions in T cells including its role in supporting Treg survival and expansion^43^ and in suppressing IL-10 expression in Th1 cells^23, 24^. Whether Bhlhe40 plays a role in Tfr cells or IL-10 upregulation is involved in Tfh cell differentiation requires further investigation. Nevertheless, our mixed bone marrow chimera experiments indicate that the role of Bhlhe40-Pou2af1 axis in regulating CXCR5 expression and thus GC-Tfh cell generation is cell intrinsic.

In conclusion, we found that the Bhlhe40-Pou2af1 axis plays an important role during Tfh maturation. Bhlhe40 represses Tfh maturation by inhibiting Pou2af1 which regulates optimal expression of the key Tfh migration factor CXCR5. Deficiency in either Bhlhe40 or Pou2af1 does not seem to affect early Tfh cell lineage commitment and Bcl6 induction. Thus, the Bhlhe40-Pou2af1 axis functions downstream of the Bcl6-Blimp1 pair and regulates pre-Tfh cells migration into geminal center to become GC-Tfh cells.

## ONLINE METHODS

### Mice

All the mouse strains are on C57BL/6 background. Wide type (WT) C57BL/6 mice were purchased from Taconic Farms. CD45.1^+^ congenic mice (line 7; Taconic), *Rag1*^-/-^ mice (line 146; Taconic) and *TCRα*^-/-^ (Line 98, Taconic) were from the National Institute of Allergy and Infectious Diseases (NIAID)–Taconic repository. *Bhlhe40*^fl/fl^ mice and *Bhlhe40*^fl/fl^-CD4Cre mice have been previously described^24^. *Bhlhe40* germline deficient allele was generated by crossing *Bhlhe40*^fl/fl^ mice with EIIa-Cre mice^44^ (JAX 003724), in which the Cre recombinase was expressed at the early mouse embryo stage resulting in germline deletion of loxP-flanked *Bhlhe40* exon 4. The following three mouse strains were generated by the CRISPR-Cas9 technology. For Bhlhe40-V5 tagged mice, the single-stranded Bhlhe40-V5 oligonucleotides (CTCTTCGGCCTTGCTCCAGGCTTTGAAGCAGATCCCTCCTTT AAACTTAGAAACCAAAGACGGCAAGCCTATCCCCAACCCTCTCCTCGGCCTCGATTC TACCTAAACTCTGGAGGGATCTCCTGCTGCCTTGCTTTCTTTCCTCCCTAATTCCAAA AACCAC), serving as the repair template, and the Bhlhe40-V5 sgRNA (ACCAAAGACTAAACTCTGGAGGG), serving as the single guide RNA (sgRNA) for targeting the *Bhlhe40* locus, were used to generate the Bhlhe40-V5 tagged mouse strain. The primer pair for identifying Bhlhe40-V5 (5’-CAAGATACCGACTCCCTTGCTTC-3’; 5’-GAATCTTCTCTGTGGGTCTGCAG-3’) was used for PCR amplification and the knock-in mutant was confirmed by DNA sequencing. The Bhlhe40-V5 tagged mutant mice were further verified by flow cytometry after staining T cells with anti-V5. To generate the *Pou2af1*^-/-^ mouse strain, two sgRNAs, Pou2af1KO-5’ (ATACCAGGGTGTTCGAGTCAAGG) and Pou2af1KO-3’ (CAACCACACCCTCTCCGTGGAGG), were used to delete exon 2-5 of the *Pou2af1* gene. The primer pair for Pou2af1 (5’-AAGCCCTGTCACTTTTAGACATAGGAGAG-3’; 5’-TGCAGACTTGGTGACATTCCGTATAAAGC-3’) was used for PCR genotyping to verify exon 2-5 deletion which was also confirmed by DNA sequencing. Two founders with exon 2-5 deletion, Pou2af1-2925-#22 (PCR product size 318 bp) and Pou2af1-2923-#20 (PCR product size 567 bp), were chosen and backcrossed with C57BL/6 for more than 6 generations to eliminate a possible off target effect of CRISPR-Cas9, and then intercrossed to generate *Pou2af1*^-/-^ mice. Pou2af1-2923-#20 was crossed with *Bhlhe40*^-/-^ to generate *Bhlhe40*^-/-^*Pou2af1*^-/-^, and other cross mice were all coming from Pou2af1-2925-#22. To generate Pou2af1-HA-tdTomato mouse strain, the single-stranded repair template was generated by inserting the DNA sequence of tdTomato followed by T2A and 3 repeats of HA tag and 10 repeats of Glycine at the ATG translation start site of the coding sequence of *Pou2af1*. The Pou2af1-HA-tdTomato sgRNA was served as the single guide RNA for targeting the *Pou2af1* locus. The primer pair Pou2af1-HA-TdTomato (5’-ACAACAACATGGCCGTCATCAAAGAGTTCA-3’; 5’-TGATGTCCAGCTTGGTGTCCACGTAGTAGTA-3’) was used for PCR and the knock-in mutant was confirmed by DNA sequencing. The Pou2af1-HA-tdTomato mutant mice were further verified by flow cytometry. *Bhlhe40*^fl/fl^ mice or *Bhlhe40*^fl/fl^-*CD4*Cre mice were crossed with Pou2af1-HA-tdTomato mice to generate Pou2af1-tdTomato reporter with or without deletion of *Bhlhe40* in CD4 T cells. Experiments were done when mice were 8–14 week of age under protocols approved by the NIAID Animal Care and Use Committee. Mice were bred and/or maintained in the NIAID specific pathogen-free animal facility.

### Mouse manipulation

In all immunization assays, mice were immunized with 20 μg AS15 peptide (AVEIHRPVPGTAPPS; synthesized by New England Peptide, Inc.) emulsified in the oil and CFA (F5881, Sigma) with volume 1:1 ratio by subcutaneous s.c. injection at two flanks along the back (50 μl/site). After 5-16 d, inguinal draining lymph nodes (dLN) or spleen were harvested to make single-cell suspension for staining or fixed for imaging. Cells were stained at 37°C for 45min with Tetramer I-Ab-AVEIHRPVPGTAPPS-APC provided by the National Institutes of Health Tetramer Core Facility.

### Bone marrow chimeras and cell transfer

For *TCRα*^-/-^ recipient mice, total bone marrow cells recovered from WT, *Pou2af1*^-/-^, *Bhlhe40*^-/-^, or *Bhlhe40*^-/-^*Pou2af1*^-/-^ mice, were mixed with *TCRα*^-/-^ bone marrow cells at a ratio of 1:5, and then adoptively transferred into sub-lethally irradiated (450 rads) *TCRα*^-/-^ mice. For *Rag1*^-/-^ recipient mice, total bone marrow cells recovered from WT CD45.1^+^ congenic mice (line 7; Taconic), were mixed with bone marrow cells from WT, *Pou2af1*^-/-^, *Bhlhe40*^-/-^, or *Bhlhe40*^-/-^*Pou2af1*^-/-^ mice at 1:1 ratio, and then adoptively transferred into sub-lethally irradiated (450 rads) *Rag1*^-/-^ mice. After 6 weeks, reconstitution was confirmed by flow cytometry, and mice were then immunized with AS15/CFA one week later. One or two weeks after immunization, cells from dLN or spleen were harvest for further analysis.

### Cell culture and purification

Naive CD4 T cells (CD4^+^CD44^lo^CD62L^hi^CD25^−^) were sorted from peripheral lymph nodes by using FACSAria (BD Biosciences Biosciences). Sorted cells were mixed with T cell depleted splenocytes^24^ with a ratio of 1:5 and then cultured under Th1 conditions (anti-CD3, 1 μg/ml; anti-CD28, 3 μg/ml; anti–IL-4, 10 μg/ml; IL-12, 10 ng/ml; and IL-2, 100 U/ml), Th2 conditions (anti-CD3, 1 μg/ml; anti-CD28, 3 μg/ml; anti–IFN-γ, 10 μg/ml; IL-12, 10 ng/ml; IL-4, 5000 U/ml; and IL-2, 100 U/ml), Th17 conditions (anti-CD3, 1 μg/ml; anti-CD28, 3 μg/ml; anti–IL-4, 10 μg/ml; anti–IFN-γ, 10 μg/ml; anti–IL-12, 10 μg/ml; TGFβ1, 1 ng/ml; IL-6, 10 ng/ml; and IL-1β, 10 ng/ml), or iTreg conditions (anti-CD3, 1 μg/ml; anti-CD28, 3 μg/ml; anti–IL-4, 10 μg/ml; anti–IFN-γ, 10 μg/ml; anti–IL-12, 10 μg/ml; TGFβ1, 5 ng/ml; and IL-2, 100 U/ml) for 4 d. Antigen specific CD4 T cells (B220^-^CD4^+^CD44^hi^AS15^+^) were FACS-sorted for three populations (PD-1^-^CXCR5^-^, PD-1^+^CXCR5^-^, PD-1^+^CXCR5^+^) from inguinal lymph nodes and used for RNA-Seq analyses. The basic culturing media is the RPMI medium 1640 (Invitrogen) supplemented with 10% FBS (Hyclone), 2 mM L-glutamine (Gibco), 1 mM sodium pyruvate (Gibco), 1X MEM non-essential amino acids (Gibco), 10 mM Hepes (Gibco), 50 μM β-mercaptoethanol (Sigma), 1X Penicillin Streptomycin (Gibco).

### Flow cytometry

Single-cell suspensions or stimulated cells were first incubated at 4 °C for 15 min with anti-CD16/32 (clone 2.4G2) for blockade of Fc receptors. Cell surface molecules were then stained with various antibodies in PBS with 3% FBS at room temperature for 30 min. For intracellular staining of transcription factors, cells were fixed and permeabilized with Foxp3 staining buffer set (00-5523-00; eBioscience) as manufacturer’s instructions. For antigen-specific (AS15^+^) cell staining, cells were pre-stained with AS15-tetramer-APC at 37 °C for 45 min before anti-CD16/32 incubation. Flow cytometry data were collected with LSRFortessa or FACSymphony (BD Biosciences) and the results were analyzed with FlowJo software (Tree Star). Antibodies were purchased from several commercial sources indicated below. Antibodies specific to mouse CD19 (eBio1D3), and PD-1 (J43) were purchased from eBioscience; antibodies specific to mouse CD45.1 (A20), CD45.2 (104), B220 (RA3-6B2) and CXCR5 (L138D7) were purchased from BioLegend; and antibodies specific to mouse CD3 (145-2C11), CD4 (GK1.5), CD44 (IM7) and Bcl6 (K112-91) were purchased from BD Biosciences. Fixable Viability Dye eFluor 506 was from eBioscience. AS15-tetramer-APC was provided by the NIH Tetramer Core Facility.

### Immunofluorescent staining

Bone marrow chimera mice were immunized with AS15/CFA, and dLNs were harvested after 7 d. Samples were treated with 1% fixation buffer (554655; BD Bioscience) at 4 °C overnight. Samples were then dehydrated in 30% sucrose and embedded in optimal cutting temperature freezing media (4585, fisher healthcare). CM1950 cryostat (Leica) and Superfrost Plus slides (VWR) were used to make sections, which were further permeabilized and blocked in PBS (0.3% Triton X-100, 1% BSA, 2% normal mouse serum and 2% normal rabbit serum) and then stained with antibodies in the same block buffer. Antibodies against IgD (11-26c.2a), CD4 (GK1.5), and CD45.2 (104) were purchased from Biolegend; antibody against CD45.1 (A20) was purchased from eBioscience; antibody against GL7 (GL7) was purchased from BD Bioscience. Fluoromount G (Southern Biotech) and Leica TCS SP8 confocal microscope were used to collect signaling from slides. Data were analyzed by Imaris software (Bitplane).

### RNA-Seq, ChIP-Seq and data analysis

For RNA-Seq analyses of Th1, Th2, Th17 and iTreg, naïve CD4 T cells from *Bhlhe40*^f/f^ or *Bhlhe40*^f/f^-CD4Cre were purified from lymph nodes and primed in vitro for 4 d. For RNA-Seq analysis of B220^-^CD4^+^CD44^hi^AS15^+^ cells, three populations (PD-1^-^CXCR5^-^, PD-1^+^CXCR5^-^, PD-1^+^CXCR5^+^) from inguinal lymph nodes were sorted from the indicated bone marrow chimera mice (WT, *Pou2af1*^-/-^, *Bhlhe40*^-/-^). The RNA-Seq was performed as previously described^45^. Briefly, the total RNAs were harvested and purified by using Qiagen’s miRNeasy micro kit (217084; Qiagen). Qiagen’s DNase set (79254; Qiagen) was used for on column DNA digestion. PolyA-tailed RNAs were purified from total RNA by using Dynabeads mRNA DIR ECT kit (61012; Ambion Life Technologies). The RNA-Seq libraries were sequenced with Illumina HiSeq system, and 50 bp reads were generated by the National Heart, Lung, and Blood Institute DNA Sequencing Core.

For ChIP-Seq analysis in Th1, Th2, Th17 and iTreg, naïve CD4 T cells from Bhlhe40-V5 tagged mice were purified from lymph nodes and primed in vitro for 4 d, and anti-V5 (R96025, Thermofisher) was used for ChIP-Seq analysis of Bhlhe40 binding in the genome. For ChIP-Seq analysis of Pou2af1 binding in the genome, splenic B cells were sorted from WT or Pou2af1-HA-TG mice, and anti-HA (ab9110, Abcam) was used for ChIP. Cells were cross-linked with 1% formaldehyde for 10 minutes at room temperature and sonicated in the shearing buffer (0.4% SDS in TE buffer) to generate 100 to 500 bp DNA fragments. 10% of total DNA was used as input, and the rest was used in the subsequent immunoprecipitation. DNA-protein complexes were pulled down with specific antibodies in the RIPA buffer overnight. DiaMag anti-mouse IgG coated magnetic beads or DiaMag protein A coated magnetic beads were added into the buffer for additional 4 hours. After washing, reverse-crosslinking and DNA purification (100 to 500 bp) were performed. Purified DNAs were then made into indexed libraries and sequenced.

For RNA-Seq and ChIP-Seq data analyses, we used the mouse reference genome mm10 and GENCODE (M21)^46^ annotations. We preprocessed all the genomic data generated in this study from raw reads to expression levels or tracks (bigWig files), following the procedures outlined in our previous studies^47, 48^. WashU Epigenome Browser^49^ was used to visualize sequencing data tracks. Specifically, we first used Cuffdiff (v2.2.1)^50^ with default parameters to identify differentially expressed genes (DEGs) between the three wild-type AS15-specific populations (1 vs. others). We used a fold change cutoff >=2 and a p-value cutoff <0.001. All DEGs from the comparison were then clustered using Cluster 3.0^51^ with the following parameters: -cg a -g 7 -m m -k 2 -r 1000. The two clusters of genes were further defined as the Tfh and Non-Tfh signature genes, which were used for the following Gene Set Enrichment Analysis (GSEA)^52^ performed by GSEApy^53^ with the subfunction of prerank and default parameters for DEGs obtained from *Pou2af1*^-/-^ cells.

### Statistics

Samples were compared with Prism 9 software (GraphPad) by a two-tailed unpaired Student’s t-test or one-way or two-way analysis of variance. A *P* value of <0.05 was considered significant. Mean ± SE; n.s., not significant, * p < 0.05, ** p < 0.01, *** p < 0.001, **** p < 0.0001.

### Accession numbers

RNA-Seq and ChIP-Seq data are available in the Gene Expression Omnibus (GEO) database (http://www.ncbi.nlm.nih.gov/gds) under the accession number GSE247137.

## ACKNOWLEDGEMENTS

We thank Ke Weng and the NIAID Flow Cytometry Section for cell sorting, NIAID animal facilities for helping maintain our mouse strains, the NIH Tetramer Core Facility for the tetramer reagents, and the NHLBI DNA Sequencing Core facility for sequencing the RNA-Seq and ChIP-Seq libraries. We thank Dechun Feng for suggestions on immunofluorescence staining, Rosalene M. Carey for helping genotyping some mouse strains, Jia Nie and Bryan Chim for helping with part of RNA-Seq analysis. This work is supported by the Division of Intramural Research of the NIAID and NHLBI.

## AUTHOR CONTRIBUTIONS

X.Z. and J.Z. conceived the project. X.Z. performed the majority of the experiments. X.C. contributed to ChIP-Seq and RNA-Seq experiments. Y.C. and G.H. performed bioinformatic analysis. C.L. helped generate knock-out and knock-in mouse strains through CRISPR-Cas9. S.G. and T.V. helped on immunofluorescence staining and signaling analysis. D.F. helped setting up antigen specific Tfh analysis. S.L. and H.C. helped analyze some bone marrow chimera mice. R.G., P.S. and K.Z. provided critical advice to the project. X.Z. and J.Z. wrote the manuscript. J.Z. supervised the project.

---

**SM Fig. 1 (related to Fig. 1).**
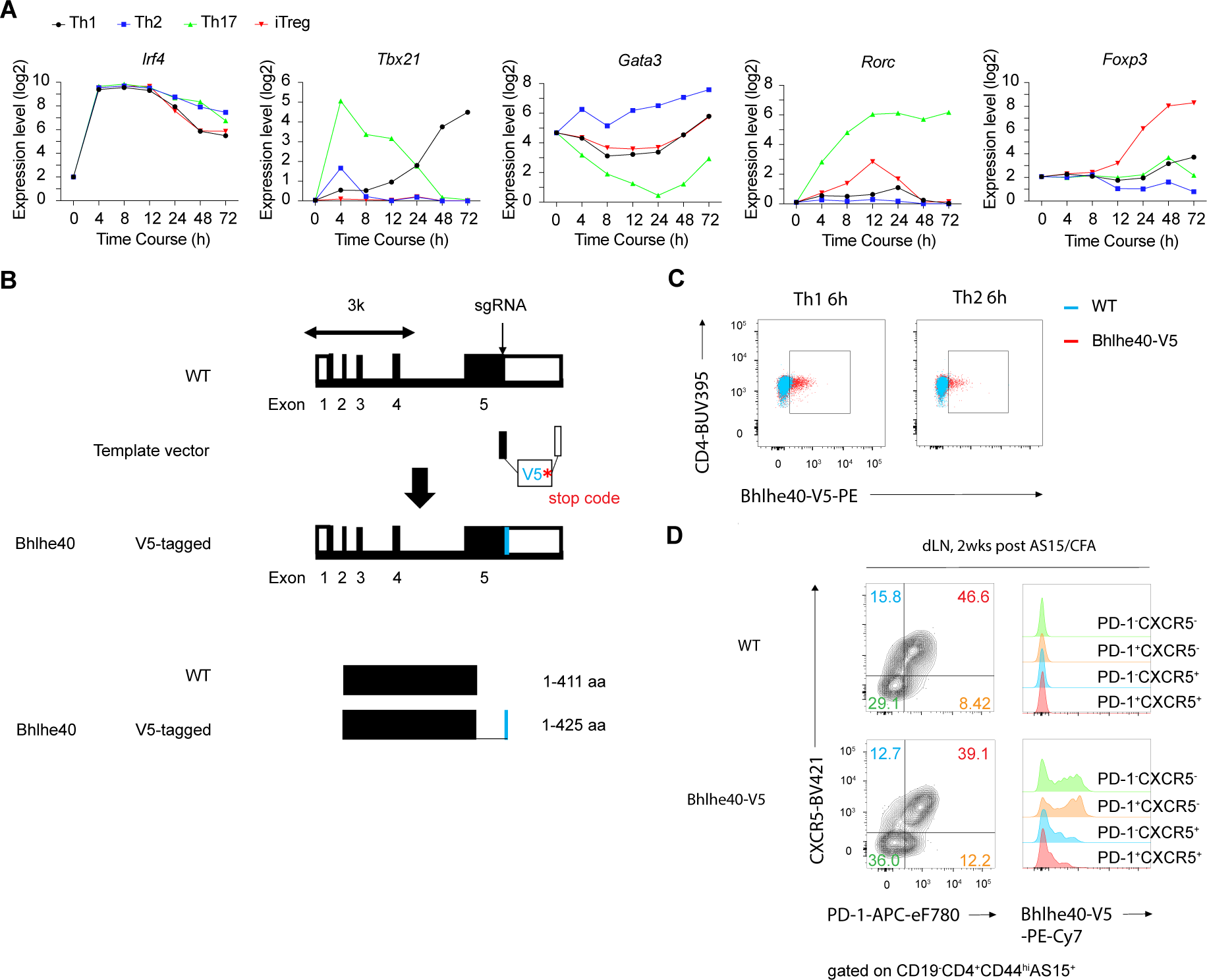
Bhlhe40 represses Pou2af1 in conventional CD4 T cells. (**A**) RNA-Seq data^33^ were re-analyzed for indicated genes expression during priming of Th1, Th2, Th17 and iTreg in vitro. (**B**) Strategy of generating Bhlhe40 V5-tagged mice. (**C**) Naïve CD4 T cells were purified from Bhlhe40-V5 mice and primed under Th1 and Th2 conditions. Bhlhe40 expression was assessed by anti-V5 staining after TCR stimulation for 6 hours. (**D**) WT and Bhlhe40-V5 mice were immunized (s.c.) with AS15/CFA for 2 weeks, and the expression of Bhlhe40 by AS15-specific effective CD4 T cells from inguinal lymph nodes divided into four populations (PD-1^-^CXCR5^-^, PD-1^+^CXCR5^-^, PD-1^-^CXCR5^+^, PD-1^+^CXCR5^+^) was assessed by flow cytometry. Data are representative of two (C and D) independent experiments.

**SM Fig.2 (related to Fig. 2).**
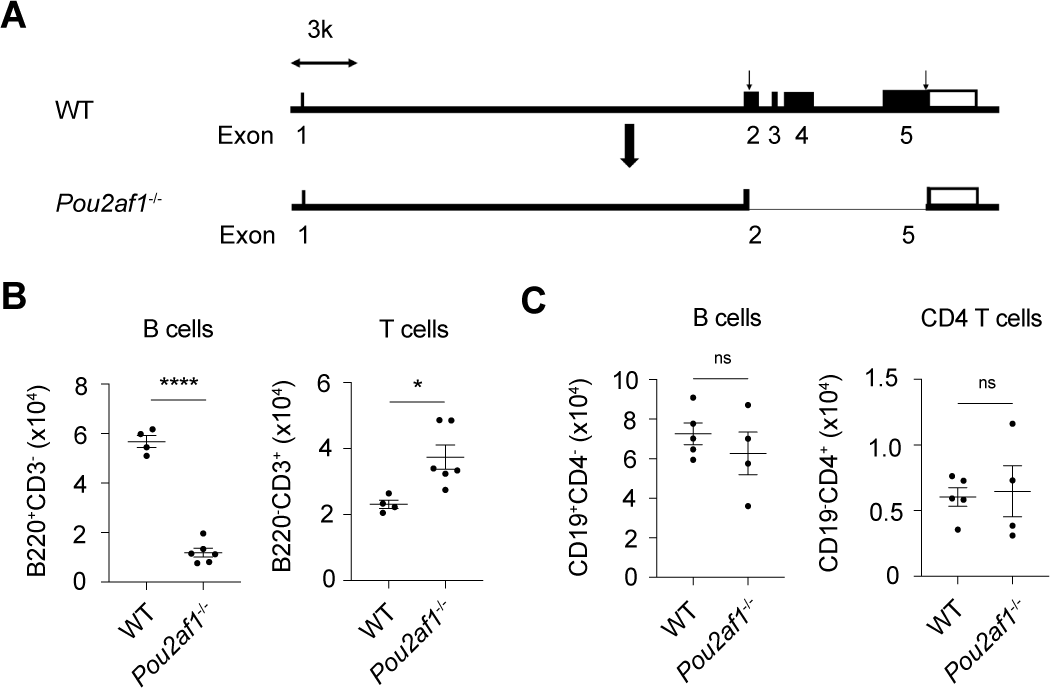
Pou2af1 is required for Tfh cell differentiation. (**A**) Strategy for generating *Pou2af1*^-/-^ mice via CRISPR-Cas9 technology. (**B**) Summary of cell number difference in B cells and T cells comparing WT (*n*=4) and *Pou2af1*^-/-^ (*n*=6) mice with the same amount of blood volume in Fig.2A. (**C**) Summary of cell number difference in B cells and CD4 T cells comparing *TCRα*^-/-^ recipients adoptively transferred with WT (*n*=5) or *Pou2af1*^-/-^ (*n*=4) with the same amount of blood volume in Fig.2E. * p < 0.05, ** p < 0.01, *** p < 0.001, **** p < 0.0001. Error bars indicate SEM. Data are representative of more than three (B) and two (C) independent experiments.

**SM Fig.3. (related to Fig. 3).**
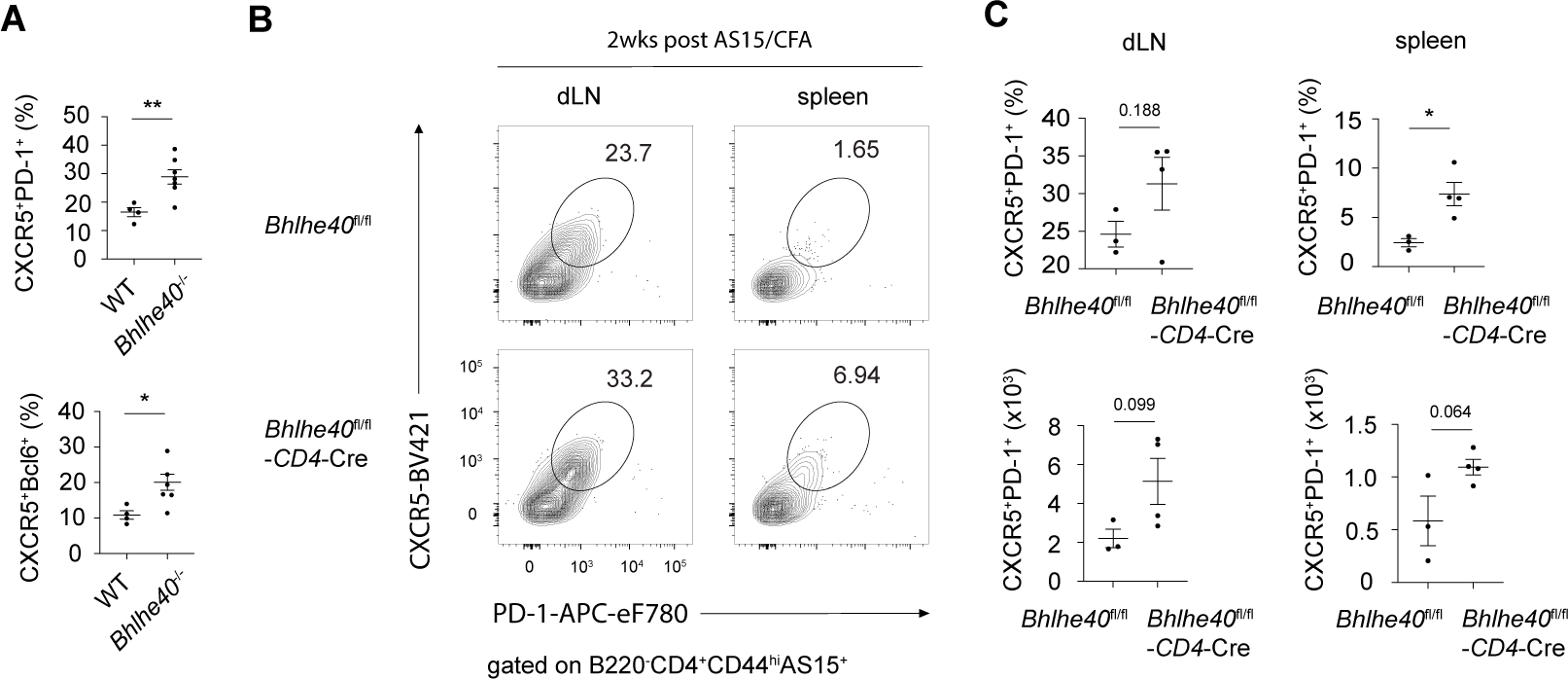
Bhlhe40 represses Tfh cell differentiation. (**A**) Summary of percentage difference in Tfh cells (PD-1^+^CXCR5^+^ or Bcl6^+^CXCR5^+^) between WT (*n*=4) and *Bhlhe40*^-/-^ (*n*=7) in the spleen in Fig. 3A. (**B**) WT (*n*=3) and *Bhlhe40*^fl/fl^-CD4Cre (*n*=4) mice were immunized (s.c.) with AS15/CFA for 2 weeks, and AS15-specific CD4 T cells from inguinal lymph nodes and spleen were analyzed for Tfh (PD-1^+^CXCR5^+^ or Bcl6^+^CXCR5^+^) cells by flow cytometry. (**C**) Summary of percentage difference in Tfh cells between WT and *Bhlhe40*^fl/fl^-CD4Cre in lymph nodes and spleen in (B). * p < 0.05, ** p < 0.01, *** p < 0.001, **** p < 0.0001. Error bars indicate SEM. Data are representative of two (A-C) independent experiments.

**SM Fig.4 (related to Fig. 4).**
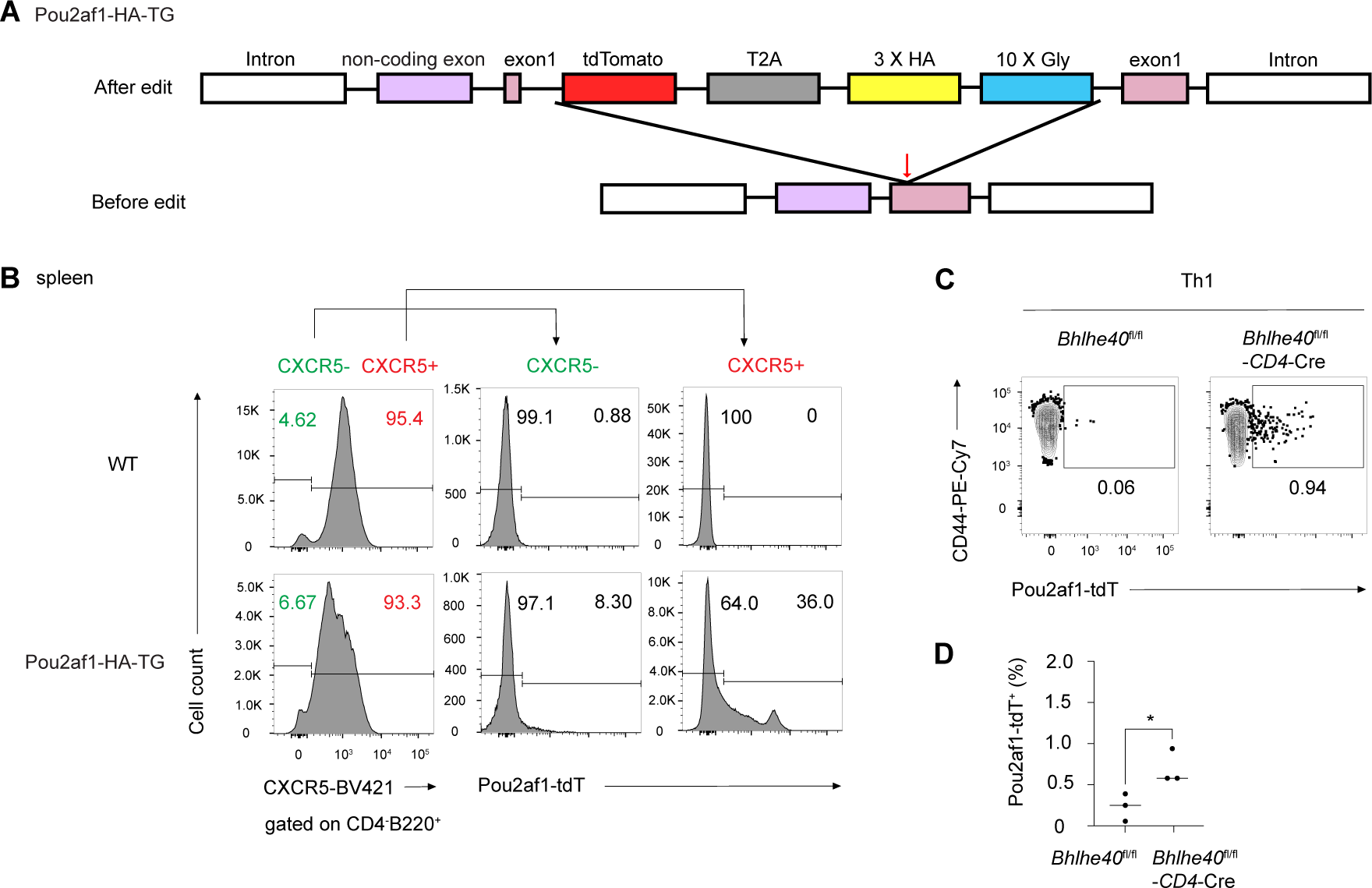
Generation of Pou2af1-HA-TG mouse strain and its usage in confirming the relationship between Pou2af1 and CXCR5 or Bhlhe40. (**A**) Strategy of generating Pou2af1-HA-TG mouse strain via CRISPR-Cas9. tdTomato (red box) and three repeats of HA (yellow box) were linked to the N-terminal of Pou2af1. (**B**) The expression of CXCR5 by B cells (CD4^-^B220^+^) from WT or Pou2af1-HA-TG mice was analyzed. CXCR5^-^ and CXCR5^+^ populations were further assessed for Pou2af1-tdTomato (Pou2af1-tdT) expression. (**C**) WT or *Bhlhe40*^fl/fl^-CD4Cre mice were crossed with Pou2af1-HA-TG mouse to generate Pou2af1-tdTomato-bearing WT (*n*=3) or *Bhlhe40*^fl/fl^-CD4Cre (*n*=3) reporter mice. Then naïve CD4 T cells were purified from such reporter mice with indicated genotypes and primed under Th1 conditions. Pou2af1 expression in Th1 cells were analyzed by flow cytometry. (**D**) Summary of percentage of Pou2af1^+^ Th1 cells in (C). * p < 0.05, ** p < 0.01, *** p < 0.001, **** p < 0.0001. Error bars indicate SEM. Data are representative of three (B) and two (C-D) independent experiments.

**SM Fig.5 (related to Fig. 5).**
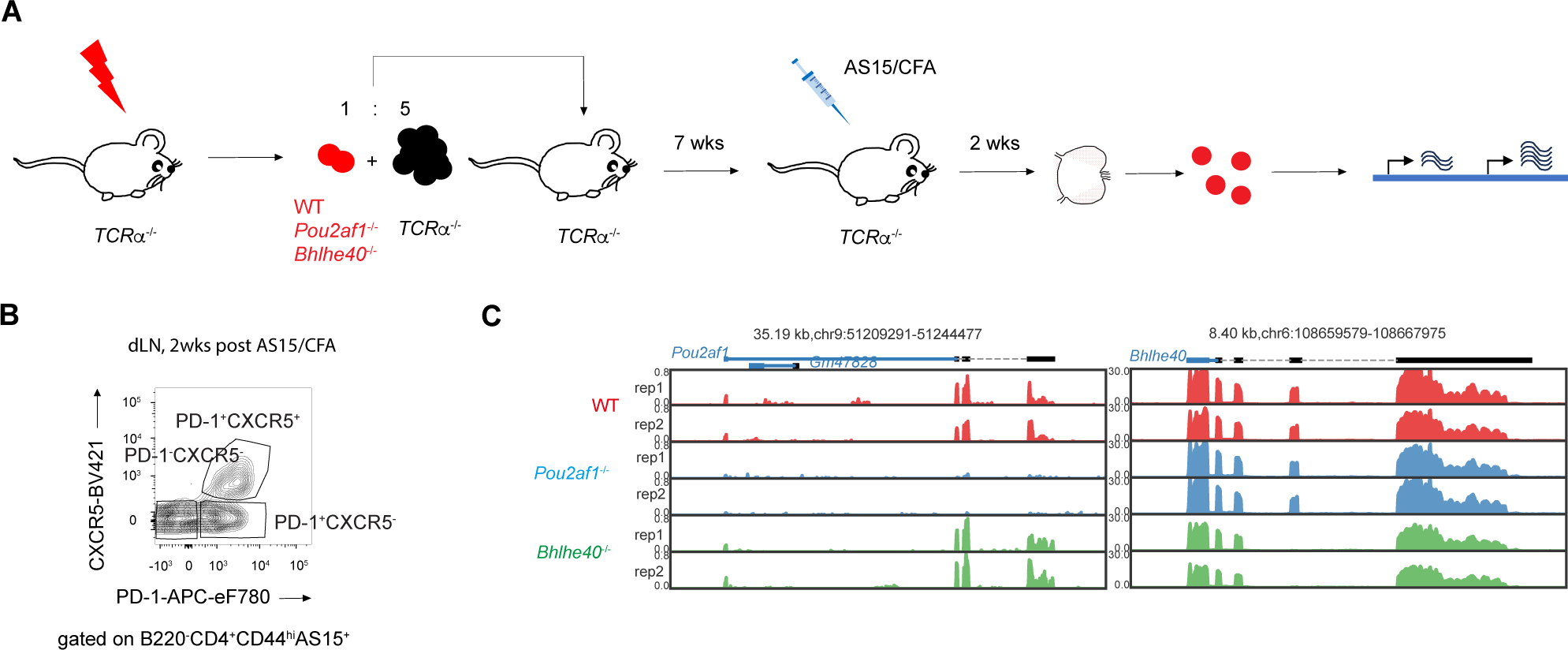
Pou2af1 and Bhlhe40 regulate Tfh differentiation in cell intrinsic manner. (**A**) Experimental procedure of immunizing bone marrow chimeras *TCRα*^-/-^ mice, received adoptively transferred bone marrow mixture of *TCRα*^-/-^ with WT, *Pou2af1*^-/-^ or *Bhlhe40*^-/-^, with AS15/CFA followed by RNA-Seq analysis. (**B**) AS15-specific CD4 T cells (B220^-^CD4^+^CD44^hi^AS15^+^) were sorted for three populations (PD-1^-^CXCR5^-^, PD-1^+^CXCR5^-^, PD-1^+^CXCR5^+^) from draining lymph nodes and used for RNA-Seq analysis. Samples are in biological duplicates. (**C**) Biological duplicates of the RNA-Seq data were shown to confirm the genotype background of the materials.

**SM Fig.6 (related to Fig. 6).**
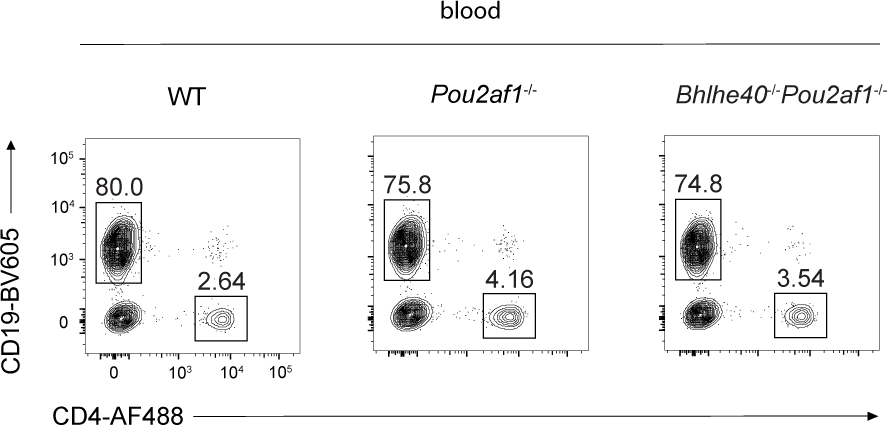
Pou2af1 functions downstream of Bhlhe40 during Tfh cell differentiation. B cells and CD4 T cells from indicated mice groups were assessed by flow cytometry in bone marrow chimeras shown in Fig. 6C.

**SM Fig.7 (related to Fig. 7).**
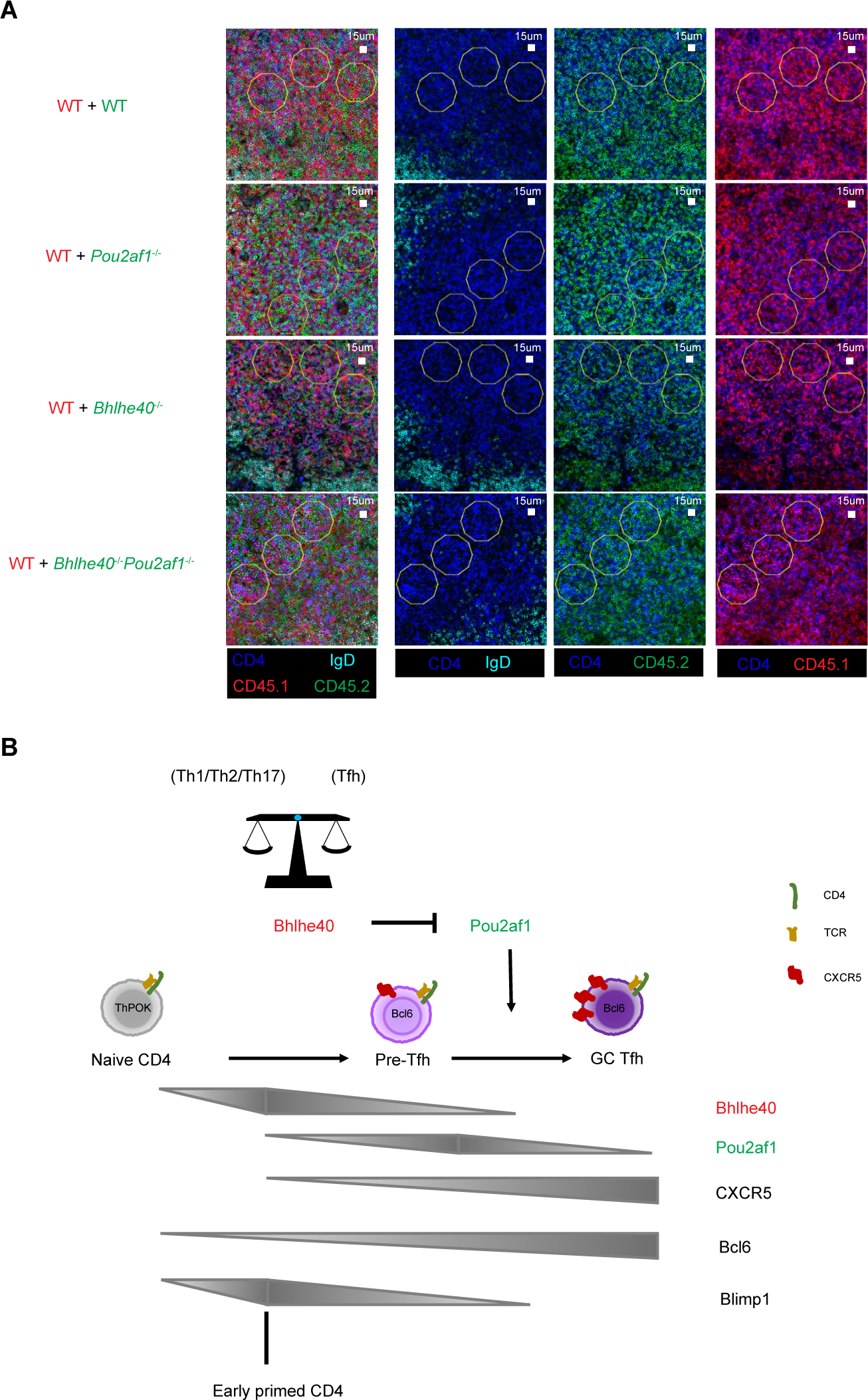
Regulation of optimal CXCR5 level through Bhlhe40-Pou2af1 axis during Tfh cell maturation. (**A**) Inguinal lymph nodes from Fig. 7A were fixed and stained for confocal image. Anti-CD4 (blue) and anti-IgD (cyan) were used to distinguish T cell area and B cell area. Three T cell areas (yellow decagon) were chosen randomly for further analysis. Anti-CD45.2 (green) as used to show different indicated genotyped adoptive cells, and anti-CD45.1 (red) was used to show WT adoptive cells from the same mouse. CD4^+^CD45.2^+^ indicated different genotype background adoptive CD4 T cells. CD4^+^CD45.1^+^ indicated WT adoptive CD4 T cells. (**B**) A model for Bhlhe40-Pou2af1 axis in regulating Tfh cell maturation by optimizing CXCR5 expression levels. Data are representative of two (A) independent experiments.

